# Long-range Temporal Correlations in the Broadband Resting state Activity of the Human Brain revealed by Neuronal Avalanches

**DOI:** 10.1101/2020.02.03.930966

**Authors:** Fabrizio Lombardi, Oren Shriki, Hans J. Herrmann, Lucilla de Arcangelis

## Abstract

Resting-state brain activity is characterized by the presence of neuronal avalanches showing absence of characteristic size. Such evidence has been interpreted in the context of criticality and associated with the normal functioning of the brain. At criticality, a crucial role is played by long-range power-law correlations. Thus, to verify the hypothesis that the brain operates close to a critical point and consequently assess deviations from criticality for diagnostic purposes, it is of primary importance to robustly and reliably characterize correlations in resting-state brain activity. Recent works focused on the analysis of narrow band electroencephalography (EEG) and magnetoencephalography (MEG) signal amplitude envelope, showing evidence of long-range temporal correlations (LRTC) in neural oscillations. However, this approach is not suitable for assessing long-range correlations in broadband resting-state cortical signals. To overcome such limitation, here we propose to characterize the correlations in the broadband brain activity through the lens of neuronal avalanches. To this end, we consider resting-state EEG and long-term MEG recordings, extract the corresponding neuronal avalanche sequences, and study their temporal correlations. We demonstrate that the broadband resting-state brain activity consistently exhibits long-range power-law correlations in both EEG and MEG recordings, with similar values of the scaling exponents. Importantly, although we observe that avalanche size distribution depends on scale parameters, scaling exponents characterizing long-range correlations are quite robust. In particular, they are independent of the temporal binning (scale of analysis), indicating that our analysis captures intrinsic characteristics of the underlying dynamics. Because neuronal avalanches constitute a fundamental feature of neural systems with universal characteristics, the proposed approach may serve as a general, systems- and experiment-independent procedure to infer the existence of underlying long-range correlations in extended neural systems, and identify pathological behaviors in the complex spatio-temporal interplay of cortical rhythms.

## 1. Introduction

The existence of long-range temporal correlations (LRTC) in resting-state cortical activity is widely recognized as a necessary ingredient for the normal functioning of the brain [1, 2, 3, 4, 5]. Recent works have shown in addition that spontaneous activity in neural systems is organized in neuronal avalanches, spatio-temporal cascades of activity that exhibit scale-invariant, power-law size distributions in both experiments [6, 7, 8, 9, 10, 11, 12, 13, 14, 15] and models [16, 17, 18, 19, 20, 21]. The concurrent emergence of LRTC and power-laws suggests that healthy neural systems may operate at criticality. This idea is further supported by the robustness of the observed scaling features, which are independent of the scale and details of the particular system, and their alterations with diseases [22]. In particular, the correlation properties of the resting state activity appear to change in the presence of brain pathology [4, 5], which indicates that reliable characterization and monitoring of correlations may represent a powerful, non-invasive diagnostic tool.

The detection and characterization of correlations in stochastic time series is a problem of wide interest, from geophysics to biology and economics, and generally is not a straightforward task, due to the presence of statistical noise, non-stationarities, and trends in the data. A number of statistical tools have been recently developed to deal with such issues and accurately assess spatio-temporal correlations. Among these, the Detrended Fluctuation Analysis (DFA) has proven especially effective in quantifying long-range power-law correlations embedded in non-stationary signals with polynomial trends and bursting dynamics [23, 24, 25, 26]. The DFA is an extension of the fluctuation analysis (FA) for determining the exponent that characterizes the scaling behavior of the fluctuations in a given signal. Unlike the FA, DFA is not affected by non-stationarities, and is therefore particularly suitable for studying LRTC in biological and physiological systems [27], as testified by its wide range of application, from the analysis of DNA sequences to cardiac dynamics [23, 28, 27, 29].

In the context of brain dynamics, DFA has been mostly used to asses LRTC in the amplitude envelope of ongoing neuronal oscillations [2, 30, 31], and corresponding alterations with diseases [5, 32, 33, 34, 35]. Specifically, *Linkenkaer-Hansen* et al [2] have shown that *α* and *β* oscillations exhibit long-range power-law correlations on time scales of hundreds of seconds during the resting-state, with scaling exponents significantly higher than the ones measured in band-pass filtered white noise [2]. Similar results, including LRTC in *δ* and *θ* waves, have been reported in several following studies [4, 36, 31], and significant changes in the scaling exponent estimated via DFA have been observed in patients affected by Alzheimer disease [5], epilepsy [33, 34], or major depressive disorders [35].

Although oscillations in specific frequency bands are usually identified with particular physiologic states, brain activity is a broadband phenomenon that likely results from a constant interplay between different brain rhythms, which may compete or temporally coexist and interact with each other [37, 38, 39]. For instance, mutual exclusion or coexistence of fast/local and slow/non-local rhythms may depend on the particular function to be performed, and requires specific cross-talk that may play a crucial part in the emergent physiologic states and associated cortical activity. Therefore, a significant piece of information useful to precisely discriminate between normal and pathological behavior, may be encoded in the complex, broadband spatio-temporal cortical dynamics.

To exploit such information, here we propose an alternative strategy to study LRTC in the broadband activity of neural systems, which combines the universality of neuronal avalanches with the effectiveness of DFA in quantifying long-range power-law correlations. Our approach consists of two steps: First, identifying avalanches and calculating their sizes; second, applying DFA to the previously obtained sequences of avalanche sizes. In the hypothesis that neural systems operate close to criticality, long-range correlations in the avalanche sequences would reflect underlying properties of neural activity embedded in the recorded signals.

We test this strategy on EEG and long-term MEG recordings (40 min) of the resting-state of the human brain. We first analyze neuronal avalanches in both data sets, and study how the size distribution depends on the scale parameters used to identify the avalanches in the recorded spatio-temporal activity. Then, we systematically investigate LRTC in the avalanche sequence, and their sensitivity to avalanche scale parameters.

## 2. Data acquisition and pre-processing

### 2.1. Resting-state MEG

Ongoing brain activity was recorded from 3 healthy female participants in the MEG core facility at the NIMH (Bethesda, MD, USA) for a duration of 40 min (eyes closed). All experiments were carried out in accordance with NIH guidelines for human subjects. The sampling rate was 600 Hz, and the data were band-pass filtered between 1 and 150 Hz. Power-line interferences were removed using a 60 Hz notch filter designed in Matlab (Mathworks). The sensor array consisted of 275 axial first-order gradiometers. Two dysfunctional sensors were removed, leaving 273 sensors in the analysis. Analysis was performed directly on the axial gradiometer waveforms.

### 2.2. Resting-state EEG

Resting-state EEG was recorded from 6 right handed healthy volunteers. Participants had no history of neurological or psychiatric diseases and had normal or corrected-to-normal vision. All gave written informed consent, and were paid for their participation. The study was approved by a local ethics committee (Ben-Gurion University) and is in accordance with the ethical standards of the Declaration of Helsinki. EEG was recorded using the g.Tec HIamp system (g.Tec, Austria) with 64 gel-based electrodes. (AgCl electrolyte gel). Electrodes were positioned according to the standard 10/20 system with linked ears reference. Impedances of all electrodes were kept below 5 kΩ. Data were pre-processed using a combination of the EEGLAB Matlab toolbox [40] routines and custom code. After high-pass filtering (cut-off 1 Hz), a customized adaptive filter was applied to suppress line-noise. This was followed by Artifact Subspace Reconstruction [41], re-referencing to the mean, and low-pass filtering (cutoff 60 Hz). Subsequently, an ICA (Independent Component Analysis) algorithm was applied to the data [42]. The resulting IC’s were evaluated automatically for artifacts by combining spatial, spectral and temporal analysis of IC’s. IC’s identified as containing ocular, muscular or cardiac artifacts were removed from data.

## 3. Neuronal Avalanches in Resting-state Brain Activity

We consider resting state-brain activity from EEG (6 healthy subjects, 62 sensors) and long-term MEG (3 healthy subjects, 273 sensors) recordings. To identify neuronal avalanches, we analyze the recorded activity over the entire sensor array, and search for spatio-temporal cascades of activity. To this end, we select positive and negative deflections at each sensor signal by applying a threshold *h* at *n* standard deviations (SD), i.e. *h* = ± *n*SD. Such procedure is illustrated in Fig. 1a for a threshold *h* = ± 3SD. In each excursion beyond the threshold, we then identify a single event at the most extreme value — i.e. maximum for positive excursions and minimum for negative excursions. This signal discretization maintains most of the strong correlations found in the continuous MEG signals recorded from different brain regions, as shown in [11].

**Figure 1:**
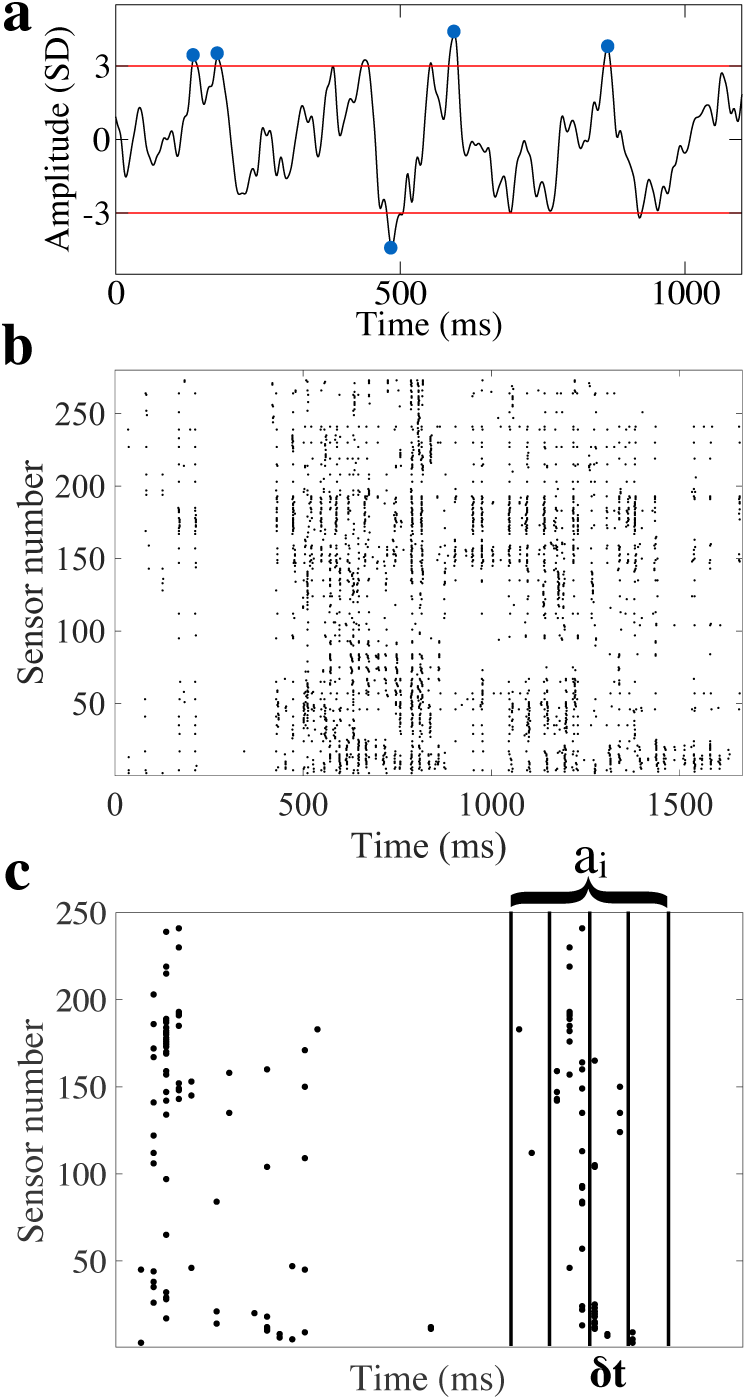
Definition of neuronal avalanches in human brain activity recordings. (a) Single sensor MEG signal of neuronal resting state activity of the human brain. The most extreme point in each excursion beyond a threshold of *h* = ± 3 SD (horizontal lines) is treated as a discrete event in the signal. In this way, the continuous signal at each sensor is mapped to a point process. (b) Raster of discrete events obtained following the procedure depicted in (a) across all MEG sensors (n = 273) over an approximately 2 s second segment of recording (single subject). (c) An avalanche *a*_*i*_ is defined as a sequence of temporal windows *δt* with at least one event in any of the sensors, preceded and followed by at least one window with no events in any of the sensors. The same procedure is used to detect avalanches in EEG recordings of resting state activity.

In Fig. 1b, we show a two second segment of the event raster extracted from the MEG of one subject after applying the discretization procedure to each of the sensor signals. We observe that events cluster in time across subgroups of sensors (Fig. 1c). A similar spatio-temporal structure can be observed in rasters of events extracted from EEG recordings (Section 2.2). The raster of events is binned at a given temporal resolutions *δt*, which is a multiple of the sampling time *T* (*T*_*meg*_: 1.67 ms; *T*_*eeg*_: 4 ms), and we define an avalanche *a*_*i*_ as a continuous sequence of time bins in which there is at least an event on any sensor, ending with at least a time bin with no events (Fig. 1c) [6]. The same identification procedure and definition of avalanches is used in both the MEG and the EEG analysis. We then define the size of an avalanche, *s*_*i*_, as the number of events in the corresponding sequence of time bins.

### 3.1. Avalanches in resting-state MEG

We first analyze the distribution *P* (*s*) of avalanche sizes from resting-state MEG recordings. We fix the time bin *δt* = 2*T*_*meg*_ = 3.3 ms, and study how *P* (*s*) depends on the value of the threshold *h* used to discretize sensor signals (Fig. 1). The distributions for a range of *h* values are shown in Fig. 2a. They generally show a power-law behavior, *P* (*s*) ∝ *s*^−*τ*^, which is independent of the threshold *h*. The power-law regime is followed by an exponential cutoff that is instead controlled by *h*, and shift from higher to smaller size *s* for increasing *h* values. Such behavior indicates that the threshold on the signal may introduce a characteristic avalanche size *s*\* ∝ *h*^−*β*^, thus suggesting that the distribution of avalanche sizes corresponding to different *h*’s may follow the general scaling form *P* (*s*) = *s*^−*τ*^ *f* (*h*^−*β*^*s*), where *τ* is the power-law scaling exponent, *f* (*h*^−*β*^*s*) is a scaling function, and *β* an exponent that expresses the dependence of the cutoff on *h*.

**Figure 2:**
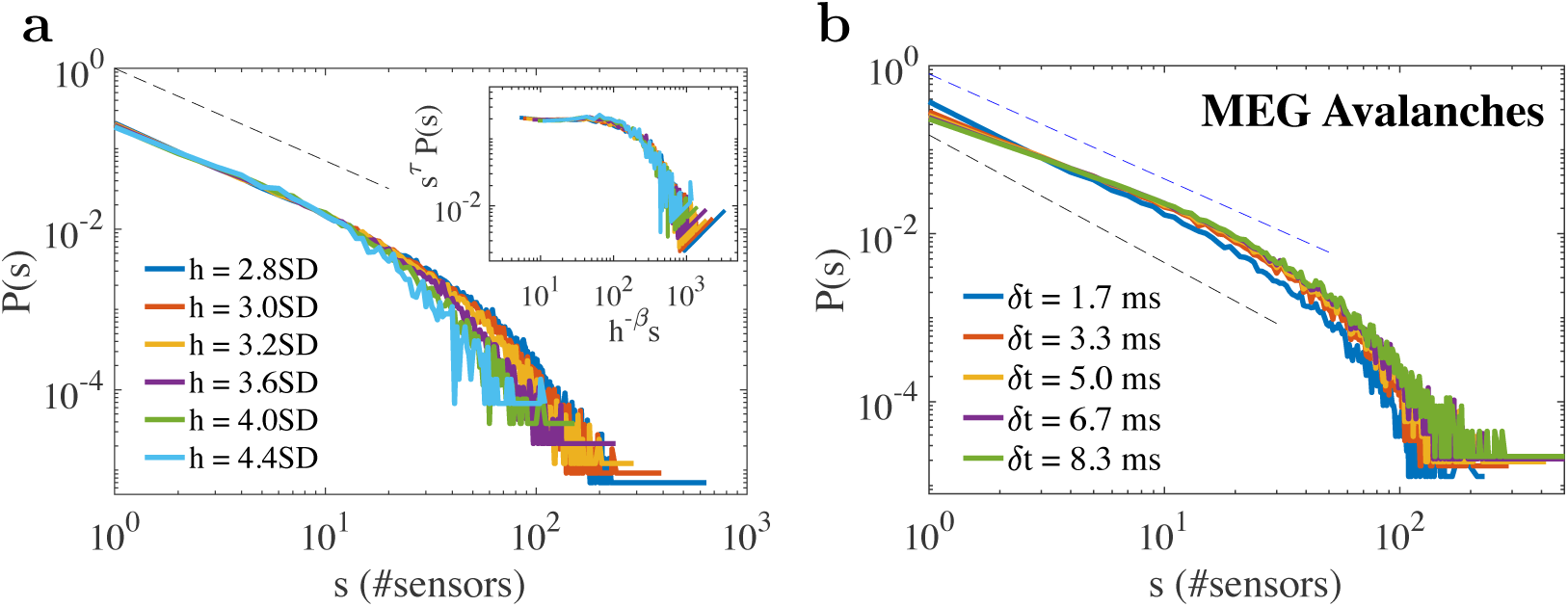
Distributions of avalanche sizes in the MEG of the resting state (pool data, 3 subjects) for different values of the threshold *h* (*δt* = 2*T*_*meg*_ = 3.3 ms), and different bin sizes *δt* (*h* = 3.0SD). The distributions generally exhibit a power-law regime followed by an exponential cutoff. (a) Distributions evaluated using different *h*’s follow a similar power-law behavior, with an exponential cutoff that is controlled by *h*. Inset: Data for different *h*’s collapse onto a single function when one plots *h*^*−β*^*s* versus *s*^*τ*^ *P* (*s*), with *τ* = 1.15 and *β* = 1.5. The dashed line in the main panel is a power-law with exponent 1.15. (b) The power-law exponent *τ* depends on the bin size *δt* used to define the avalanches. *δt* ranges between 1*T*_*meg*_ = 1.67 ms to 5*T*_*meg*_ = 8.3 ms. For increasing values of *δt*, the probability for large (small) avalanches increases (decreases) and, correspondingly, *τ* tends to decrease. The black and blue dashed lines are power-laws with exponents 1.52 and 1.25, respectively.

We then use such assumption to determine the scaling exponents *τ* and *β* by plotting *s*^*τ*^ *P* (*s*) versus *h*^−*β*^*s* for the different *h* values (inset of Fig. 2a). We found that the distributions collapse onto a single curve *f* (*h*^−*β*^*s*) for *τ* = 1.15 and *β* = 1.5. The value of the exponent *τ* obtained from the data collapse is slightly smaller than those reported in previous works, where *τ* has been estimated via maximum likelihood over the entire range of avalanche sizes [11]. The observed difference could be due to the deviation from an optimal data collapse along the scaling regime (inset of Fig. 2a). Indeed, least square or maximum likelihood estimates of the power-law exponent *τ* provide values that are slightly higher and closer to those reported in other experiments [11], as we shall discuss hereinafter.

Next, we keep the threshold *h* fixed (*h* = 3SD) and examine the size distribution varying the bin size *δt*, which controls the temporal scale of the analysis. We observe that the size distribution follows a similar functional behavior for all *δt* values, namely a power-law followed by an exponential cutoff (Fig.2b). However, the power-law exponent *τ* depends on *δt*, and tends to decrease for increasing *δt* values. Indeed, a larger bin size *δt* tends to merge small events into larger ones, leading to an increase in the probability to observe larger avalanches and thus to a smaller scaling exponent *τ*. We find that the exponent goes from *τ* = 1.52 ± 0.02 for *δt* = *T*_*meg*_ = 1.67 ms, to *τ* = 1.25 ± 0.02 for *δt* = 5*T*_*meg*_ = 8.3 ms (exponents are estimated over the size range [1,40] for *δt* = 1.67 ms, [1,50] otherwise, i.e. excluding the exponential cutoff) (Fig.2b). These values are in line with previous estimates on resting-state MEG data [11].

### 3.2. Avalanches in resting-state EEG

The analysis of the size distribution *P* (*s*) gives similar results for the neuronal avalanches in the resting EEG. The distribution *P* (*s*) shows a power-law behavior with an exponential cutoff, independently of the particular values of the bin size and threshold used to identify the avalanches (Fig. 3). However, we find that its scaling exponent *τ* is slightly larger than in MEG recordings. Moreover, unlike our observations in the resting MEG, *τ* appears to be more sensitive to the choice of data discretization thresholds *h* (Fig. 3a). Indeed, data do not collapse in the full range of threshold values, as found in the resting MEG, but only asymptotically for *h >* 3.4SD. Our analysis shows that *τ* is close to 1.5 (1.53 ± 0.04) for *h* = 2.8SD and increases for increasing *h* values, while the exponential cutoff shift to smaller size *s* (Fig. 3a). For *h >* 3.4SD, the distributions approximately collapse onto a unique function *f* (*h*^−*β*^*s*) satisfying the relation *P* (*s*) = *s*^−*τ*^ *f* (*h*^−*β*^*s*) (Section 3), with *τ* = 1.85 and *β* = 1.5 (inset of Fig. 3a).

**Figure 3:**
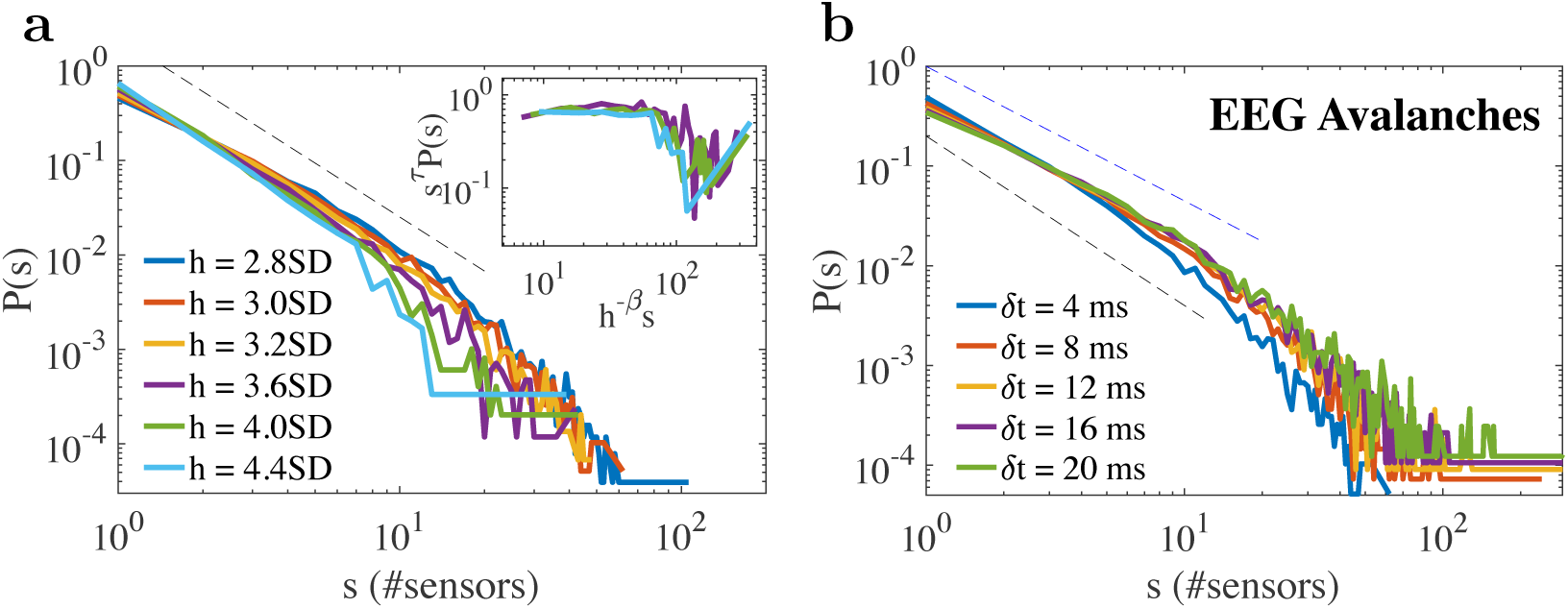
Distributions of avalanche sizes in the EEG of the resting state (pool data, 6 subjects) for (a) different values of the threshold *h* (*δt* = 2*T*_*eeg*_ = 8 ms), and (b) different bin sizes *δt* (*h* = 3.0SD). The distributions generally exhibit a power-law regime followed by an exponential cutoff, as observed in the MEG (Fig. 2). (a) The power-law exponent *τ* depends on the threshold *h*, in particular for small *h*’s, and tends to increase for increasing threshold values. The exponential cutoff is controlled by *h*, and with increasing *h* values, shifts to smaller avalanche sizes *s*. Inset: Data for *h >* 3.4 collapse onto a single function when one plots *h*^*−β*^*s* versus *s*^*τ*^ *P* (*s*), with *τ* = 1.85 and *β* = 1.5. The dashed line in the main panel is a power-law with exponent 1.85 (b) The power-law exponent *τ* also depends on the bin size *δt* used to define the avalanches, as already observed for MEG avalanches. *δt* ranges between 1*T*_*eeg*_ = 4 ms to 5*T*_*eeg*_ = 20 ms. For increasing values of *δt*, the probability for large (small) avalanches increases (decreases) and, correspondingly, the exponent *τ* tends to decrease. The black and blue dashed lines are power-laws with exponents 1.7 and 1.36, respectively.

We also analyze how the scaling exponent *τ* depends on the bin size *δt* (Fig. 3b). We find that *τ* tends to decrease for increasing bin sizes, as already observed for avalanches in the resting-state MEG (Fig. 2b). For a threshold *h* = 3SD, the power-law exponent goes from *τ* = 1.70 ± 0.04 for *δt* = 1*T*_*eeg*_ = 4 ms, to *τ* = 1.36 ± 0.02 for *δt* = 5*T*_*eeg*_ = 20 ms. Exponents are estimated considering sizes in the range *s* < 20. The reported values are slightly higher than those found in the resting MEG for the same range of parameters, namely *h* = 3SD and *δt* comprised between one and five times the sampling interval (Fig. 3).

## 4. Long-range correlations in resting-state brain activity

We have shown that resting-state brain activity is consistently organized in neuronal avalanches, distributed clusters of neuronal activity whose sizes are power-law distributed. To identify avalanches in the MEG and EEG recordings we used a discretization procedure first, and then binned the resulting discrete raster of events. Although the power-law exponent of the size distribution slightly depends on the parameters involved in the identification procedure, the general functional behavior of the distribution, i.e. power-law with an exponential cutoff, does not, indicating the fundamental character of the spatio-temporal organization captured by avalanche analysis [6]. Moreover, the signal discretization procedure illustrated in Section 3 retains the correlations present in the signal [11], and therefore makes avalanches suitable to investigate the underlying properties of resting state activity.

Building upon this fundamental results, in the following we consider the sequences of avalanches previously obtained and investigate LRTC using DFA. The DFA [23] is designed to quantify long-range power-law correlations in non-stationary signals with polynomial trends and intermittent bursting dynamics [24, 25, 26]. It consists of the following steps: (i) Given a sequence of *N* avalanche sizes *s*_*i*_(*i* = 1, …, *N*) calculate the integrated signal 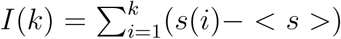, where < *s* > is the mean of *s*_*i*_ and *k* = 1, …, *N*; (ii) Divide the integrated signal *I*(*k*) into boxes of equal length *n* and, in each box, fit *I*(*k*) with a first order polynomial *I*_*n*_(*k*), which represents the trend in that box; (iii) For each *n*, detrend *I*(*k*) by subtracting the local trend, *I*_*n*_(*k*), in each box and calculate the root-mean-square (r.m.s.) fluctuation 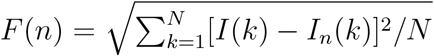; (iv) Repeat this calculation over a broad range of box lengths *n* and obtain a functional relation between *F* (*n*) and *n*. For a power-law correlated time series, the average r.m.s. fluctuation function *F* (*n*) and the box size *n* are connected by a power-law relation, that is *F* (*n*) ∝ *n*^*α*^. The exponent *α* is a parameter which quantifies the long-range power-law correlation properties of the signal. Values of *α <* 0.5 indicate the presence of anti-correlations in the sequence *s*_*i*_, *α* = 0.5 absence of correlations, and *α >* 0.5 indicates the presence of positive correlations in *s*_*i*_.

In Fig. 4 we show the fluctuation function *F* (*n*) as function of *n* for avalanche sequences from MEG recordings. We observe that *F* (*n*) ∝ *n*^*α*^, with the scaling exponent *α* consistently larger than 0.5, indicating the presence of long-range power-law correlations among avalanches, and thus in the resting-state brain activity. Importantly, we find that, for a given fixed value of the threshold *h, α* is very robust and independent of the bin size *δt*. In Fig. 4a we show the fluctuation function for avalanche sequences with *h* = 3SD, for which we find *α* ≈ 0.72 for all bin sizes (*α* = 0.716 ± 0.004 for *δt* = 1.67 ms; *α* = 0.7206 ± 0.006 for *δt* = 8.3 ms). This indicates that, although the temporal binning influences the scaling behavior of the avalanche size distribution, it does not alter the property of the correlations.

**Figure 4:**
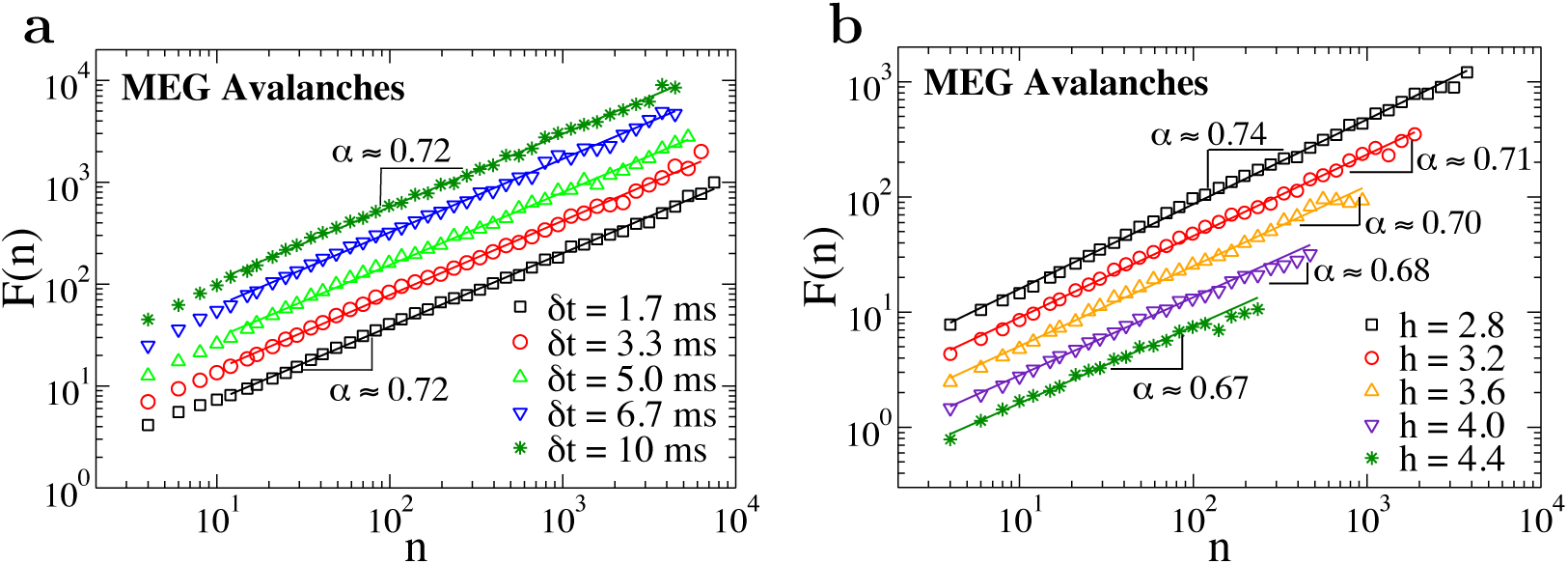
Detrended fluctuation analysis for sequences of MEG avalanche sizes extracted using different values of the parameters *h* (a) and *δt* (b). The rms fluctuation function *F* (*n*) is obtained averaging over all subjects. Loglog plots of *F* (*n*) versus the time scale of analysis *n*, where *n* is the number of consecutive avalanche sizes, show power-law relations *F* (*n*) ∝ *n*^*α*^. (a) The scaling exponents are significantly larger than 0.5, which indicates presence of positive (persistent) long-range correlations in avalanche sizes, and are independent of the bin size *δt* chosen to identify neuronal avalanches. The threshold *h* is fixed and equal 3SD. (b) The exponent *α* is slightly dependent of the threshold *h*, and fluctuates in the interval 0.7 ± 0.03 (b). *δt* = 3.3 ms for all curves.

Next, we examine the behavior of the fluctuation function for avalanche sequences obtained for a fixed bin size *δt* = 3.3 ms and different thresholds *h*. We find that the exponent *α* is always significantly larger than 0.5, indicating that the discretization procedure does not affect the nature of correlations, and that LRTC in avalanche sequence reflect the underlying properties of the signal. The scaling exponent *α* slightly depends on *h*, and decreases for increasing *h* values (*α* = 0.7360 ± 0.0056 for *h* = 2.8SD; *α* = 0.6694 ± 0.0114 for *h* = 4.4 SD). Such behavior is reminiscent of what is observed in other natural stochastic processes, as earthquake triggering, where large events are Poissonian but trigger a number of smaller and correlated in time successive events [43]. This suggests that larger values of *h* may tend to select extreme events which are less correlated than smaller ones. However, for the range of thresholds analyzed, we find that the scaling exponent is always significantly larger than 0.5, namely *α* ≈ 0.70 ± 0.03. Moreover, we observe that differences in the exponent values tend to be less relevant for *h >* 3.2SD, (*α* = 0.6950 ± 0.0100 for *h* = 3.6SD; *α* = 0.6773 ± 0.008 for *h* = 4.0SD; *α* = 0.6694 ± 0.0114 for *h* = 4.4SD) (Fig. 4b).

The same analysis performed on sequences of avalanches from EEG recordings confirms the presence of LRTC in broadband resting-state brain activity (Fig.5). In particular, we observe that the scaling exponent *α* is significantly larger than 0.5 and, for fixed values of the threshold *h*, is independent of the bin size used to detect neuronal avalanches (*α* = 0.6651 ± 0.0073 for *δt* = 4 ms; *α* = 0.6467 ± 0.0088 for *δt* = 20 ms) (Fig.5a). Furthermore, the scaling behavior of the fluctuation function *F* (*n*) appears to be also independent of the threshold *h*, the power-law exponent *α* being consistently in the range [0.66, 0.68] (*α* = 0.6806 ± 0.0061 for *h* = 2.6SD; *α* = 0.6652 ± 0.0073 for *h* = 3.0SD; *α* = 0.6820 ± 0.0069 for *h* = 3.4SD; *α* = 0.6721 ± 0.0183 for *h* = 4.0SD) (Fig.5b). We notice that the scaling exponent *α* tends to be smaller for EEG than MEG recordings, while EEG avalanches generally exhibited a larger power-law exponent *τ* in the size distribution (Section 3). This seems to indicate that *α* and *τ* are anti-correlated. However, further investigations are needed to confirm such preliminary evidence.

**Figure 5:**
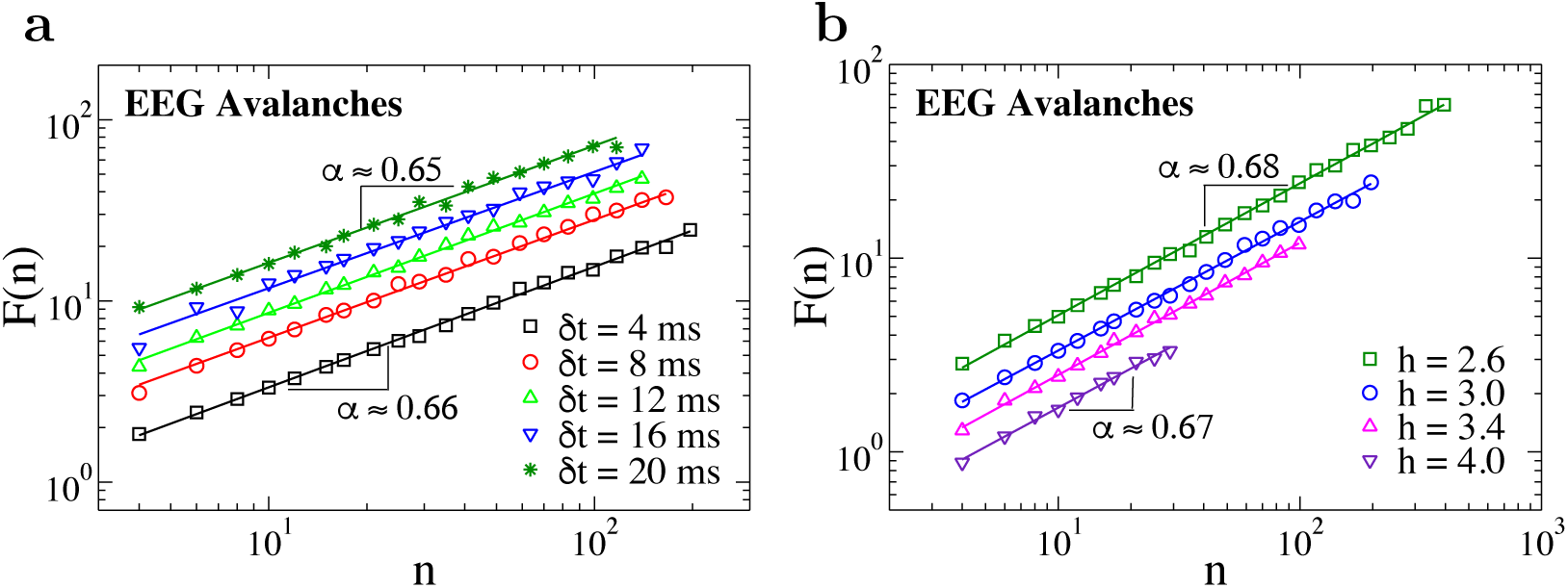
Detrended fluctuation analysis for sequences of EEG avalanche sizes extracted using different values of the parameters *h* (*δt* = 4 ms) (a) and *δt* (*h* = 3SD) (b). The rms fluctuation function *F* (*n*) is obtained averaging over all subjects. Loglog plots of *F* (*n*) versus the time scale of analysis *n*, show power-law relations *F* (*n*) ∝ *n*^*α*^, as observed for MEG avalanches (Fig. 4). The scaling exponent *α* is significantly larger than 0.5 for all values of the parameters, indicating presence of positive (persistent) long-range correlations also in EEG avalanche sizes. *α* is close to 0.7, as in the resting MEG (Fig. 4), and is independent of both the bin size *δt* and threshold *h* chosen to identify neuronal avalanches..

The results of the DFA on the sequences of neuronal avalanches consistently show the presence of LRTC both in broadband MEG and EEG recordings of the resting-state brain activity, with a scaling exponent *α* ≈ 0.7 for the fluctuation function *F* (*n*). Crucially, such an exponent is independent of the parameter *δt*, which controls the temporal binning used to identify avalanche sequences in the recordings, and thus the scale of observation of the spatio-temporal dynamics. This indicates that coarse-graining does not affect correlation properties, namely they are scale invariant as in systems at criticality, and thus the presented results capture intrinsic features of the broadband resting-state activity in the human brain.

## 5. Conclusions

We have shown that the broadband resting brain activity from MEG and EEG can be effectively described as sequences of spatio-temporal clusters of events, i.e. neuronal avalanches, and that such characterization can be used to efficiently extract relevant information about the underlying dynamics. In both our MEG and EEG recordings, neuronal avalanches exhibited similar features, with a power-law behavior in the size distribution characterized by a similar value of the scaling exponent *τ*. We first studied the dependence of this exponent on the scale parameters of the avalanche identification procedure, namely the discretization threshold *h* and the bin size *δt*, and showed that, both in the MEG and EEG recordings, its value decreases for increasing bin sizes. This effect is due to the merging of smaller avalanches into larger ones, as pointed out in [6]. On the other hand, our results showed that the power-law exponent is weakly dependent of the threshold *h*, and distributions approximately collapse onto a unique function for a range of *h* values.

We then used the avalanche representation of resting-state brain activity — i.e. the sequence of avalanches extracted from the recordings — to assess long-range temporal correlations by means of DFA. Our analysis consistently showed the presence of power-law LRTC in both MEG and EEG recordings, with exponent *α* ≈ 0.7. LRTC in EEG recordings tend to be characterized by scaling exponents slightly smaller than those measured in MEG data. On the contrary, we observed that EEG avalanche size distributions generally exhibit a larger power-law exponent *τ* as compared to MEG avalanches. These results may represent a preliminary indication that *α* and *τ* are anti-correlated, which needs further validation.

Remarkably, despite the observed dependence of avalanche size distribution on parameters, the exponent characterizing LRTC is quite robust. In particular, such exponent is independent of the temporal binning, indicating that correlation features are independent of the scale of analysis, and therefore that our approach captures intrinsic characteristics of the underlying dynamics. Furthermore, the signal discretization involved in the identification of avalanches does not crucially influence the correlations present across sensor signals [11], as demonstrated by the weak (MEG) or no (EEG) dependence of the scaling exponent *α* on the threshold *h*.

Overall these results show that the proposed approach is a valuable, effective, and broadly applicable strategy to study LRTC in neural systems. Unlike previous studies on LRTC in brain activity [2, 4, 5, 31], our approach does not focus on specific neural oscillations, but considers the intrinsic broadband nature of the resting-state brain activity and its complex spatio-temporal organization. The reported values of the LRTC scaling exponent are in the range of those observed for neural oscillations [2, 4, 5, 31]. Specifically, they are very close to the exponents characterizing oscillations in the *α* band (≈ 0.7) [2, 5, 31], suggesting that broadband correlation properties are mostly controlled by the dominant frequency band, e.g. alpha in the resting state. Importantly, this approach overcomes the limitations connected to the use of Hilbert transform in previous studies on neural oscillations, and is suitable to be extended to narrow band avalanches.

The study of temporal correlations in neuronal avalanches generally constitutes a powerful tool to understand emergent collective neural dynamics and assess presence of underlying criticality. Indeed, power-law behavior in avalanche size and duration distributions do not necessarily imply criticality, and could also arise, for instance, from the superposition of independent Poisson processes with a common temporal envelope [44]. However, in such a case the system would not show long-range correlations. Thus, our analysis of LRTC becomes crucial to properly identify genuine critical behavior in neural systems. Previous studies on cortical dynamics at a much smaller scale have shown that the temporal organization of avalanches of different sizes is far from being random, and carries a wealth of information about the deep level of self-organization in neural activity [45, 46]. For instance, detailed studies [47, 45, 48] in terms of conditional probabilities have shown the importance of the E/I balance in the emergent neural dynamics, and its connection with the up/down state dynamics: During the up-states the sizes of successive avalanches are correlated, and typically an avalanche tend to be followed by a close-in-time smaller avalanche; conversely, avalanches separated by a longer time delay, i.e. of the order of the duration of a down-state, tend to have a reversed relationship, with the second avalanche larger than the first one. The outcome of this emergent organization of avalanches in time is the observation of brain modes in the global activity [49]: Large avalanches tend to occur at slow frequency (*θ* rhythm), whereas smaller avalanches occur at higher frequency (*β/γ* frequency). In this scenario, the observation of nested oscillations is fully coherent with scale-free scaling properties of neuronal avalanches [9]. On the other hand, altering the E/I ratio disrupts such relation between avalanche sizes, and the activity can be potentially abnormal with growing avalanche sizes also at short timescales [45, 48].

The analysis presented here aims at exploiting the universal character of neuronal avalanches and the information they carry about the underlying dynamics, and provides a general approach for investigating LRTC in neural systems. The robustness of the observed scaling behavior in the rms fluctuation function and the versatility of the proposed scheme — which is suitable for both broad- and narrow-band signals —, make this approach a valuable tool to investigate alterations of LRTC in neurological diseases. Moreover, the prospect of establishing a specific relationship between avalanche size and broadband LRTC scaling exponents may greatly simplify the problem of characterizing neuronal avalanche dynamics across systems and conditions, and reliably assessing significant differences among them.

## 6. Acknowledgements

LdA would like to acknowledge the financial support from MIUR-PRIN2017 WZFTZP and VALERE:VAnviteLli pEr la RicErca 2019. FL acknowledges support from the European Union’s Horizon 2020 research and innovation programme under the Marie Sklodowska-Curie Grant Agreement No. 754411. HJH would like to thank the Agencies CAPES and FUNCAP for financial support.

